# Electrophysiological lag threads reveal a temporal hierarchy of the human cortex

**DOI:** 10.64898/2026.05.27.728161

**Authors:** Paul Hege, Janet Giehl, Markus Siegel

## Abstract

The human brain is spatially organized along functional gradients, but the temporal organization of these gradients remains poorly understood. In EEG and MEG, this question has been difficult to address because volume conduction obscures temporal precedence. Here, we introduce lag-specific orthogonalization, an extension of pairwise orthogonalization that allows us to estimate non-zero-lag amplitude-envelope correlations. Applying this approach to large-scale resting-state MEG, we identify electrophysiological lag threads: reproducible maps of temporal ordering that are most coherent in the alpha and beta bands and organize activity from occipital and sensorimotor regions toward frontal and temporal association cortex. These fast electrophysiological lag threads share subject-specific spatial structure with much slower fMRI lag threads, supporting a neural rather than vascular origin of fMRI lag structure. Finally, we show that cortical lag threads are not fixed anatomical constraints. They covary with age and handedness and are selectively reconfigured during cognitive task performance. Together, these findings reveal a frequency-specific electrophysiological temporal hierarchy of the human cortex that links fast neural dynamics to slower hemodynamic propagation structure.

## Introduction

The human functional connectome is spatially organized along macroscale gradients, with a principal axis extending from unimodal sensory and motor cortex to transmodal association areas, including the default mode network ^1,2^. This spatial hierarchy is thought to constrain how cortical regions process and integrate information. It changes across the lifespan ^3^, and is disrupted in several neurological and psychiatric conditions, including autism spectrum disorder, schizophrenia, and major depressive disorder ^4–7^. However, while the spatial topography of these networks is well characterized, their temporal organization remains much less clear. In particular, it remains unclear when, and in which sequence, activity emerges across cortical regions.

Resting-state fMRI has provided evidence that spontaneous cortical activity is not simultaneous, but unfolds in structured and reproducible sequences across the cortex, termed lag threads ^8,9^. These sequences broadly follow the principal spatial gradient, with unimodal regions tending to lead transmodal regions. However, fMRI measures neural activity only indirectly through hemodynamic responses. Thus, regional differences in neurovascular coupling could produce apparent temporal ordering without reflecting neural dynamics ^10^. Direct electrophysiological measurements are thus needed to test whether cortical lag structure has a neural basis and how it is expressed on the faster timescale of population activity.

MEG provides the temporal resolution required to study fast cortical dynamics. Previous studies have investigated lagged interactions of cortical activity using source-reconstructed MEG^11,12^ and other direct measures of electrophysiological activity.^13,14^ However, non-zero lag amplitude correlations in MEG have been fundamentally limited by volume conduction and signal mixing ^15–18^. These effects generate strong artifactual zero-lag correlations. Existing orthogonalization methods address this problem at zero lag, but leave substantial residual artifacts at non-zero lags. Consequently, the temporal ordering of amplitude fluctuations across cortical regions has remained difficult to access with MEG.

Here, we extend pairwise, time- and frequency-specific orthogonalization ^18^ to arbitrary temporal lags by introducing a lag-dependent phase precession in the complex wavelet domain. This lag-specific orthogonalization removes signal mixing artifacts from non-zero-lag amplitude envelope correlations and thereby allows us to map temporal precedence in source-reconstructed MEG. Applying this method to large-scale resting-state MEG data, we identify frequency-specific electrophysiological lag threads: reproducible maps of cortical temporal ordering that are most coherent in the alpha and beta bands and organize activity from posterior and sensorimotor regions toward frontal and temporal association cortex. In matched subjects, these fast electrophysiological lag threads partially share subject-specific spatial structure with much slower fMRI lag threads, supporting a neural contribution to hemodynamic lag structure. Finally, we show that lag threads covary with handedness and age within young adulthood and are selectively reconfigured during cognitive task performance. Together, these findings reveal a frequency-specific electrophysiological temporal hierarchy of the human cortex.

## Results

### Lag-specific orthogonalization eliminates artifactual correlations

Measures of neural covariation derived from electromagnetic recordings are strongly affected by volume conduction and signal mixing. In M/EEG, imperfections in the forward model, re-referencing, and the ill-posed nature of source reconstruction mean that even source-level estimates contain artifactual correlations ^15–18^. At zero lag, established methods can remove these artifacts ^18,19^. However, substantial spurious correlations remain at non-zero lags. To illustrate this, we simulated two independent white-noise signals with controlled instantaneous mixing. Without orthogonalization, this mixing produced a strong spurious cross-correlation at zero lag (Fig. 1a, red). Standard zero-lag orthogonalization ^18^ removed the zero-lag artifact, but left an oscillatory artifact at non-zero lags with a periodicity of twice the base frequency. This artifact results from the fact that real-valued wavelet filters are not orthogonal at arbitrary temporal shifts (Fig. 1a–c, blue).

**Fig 1.**
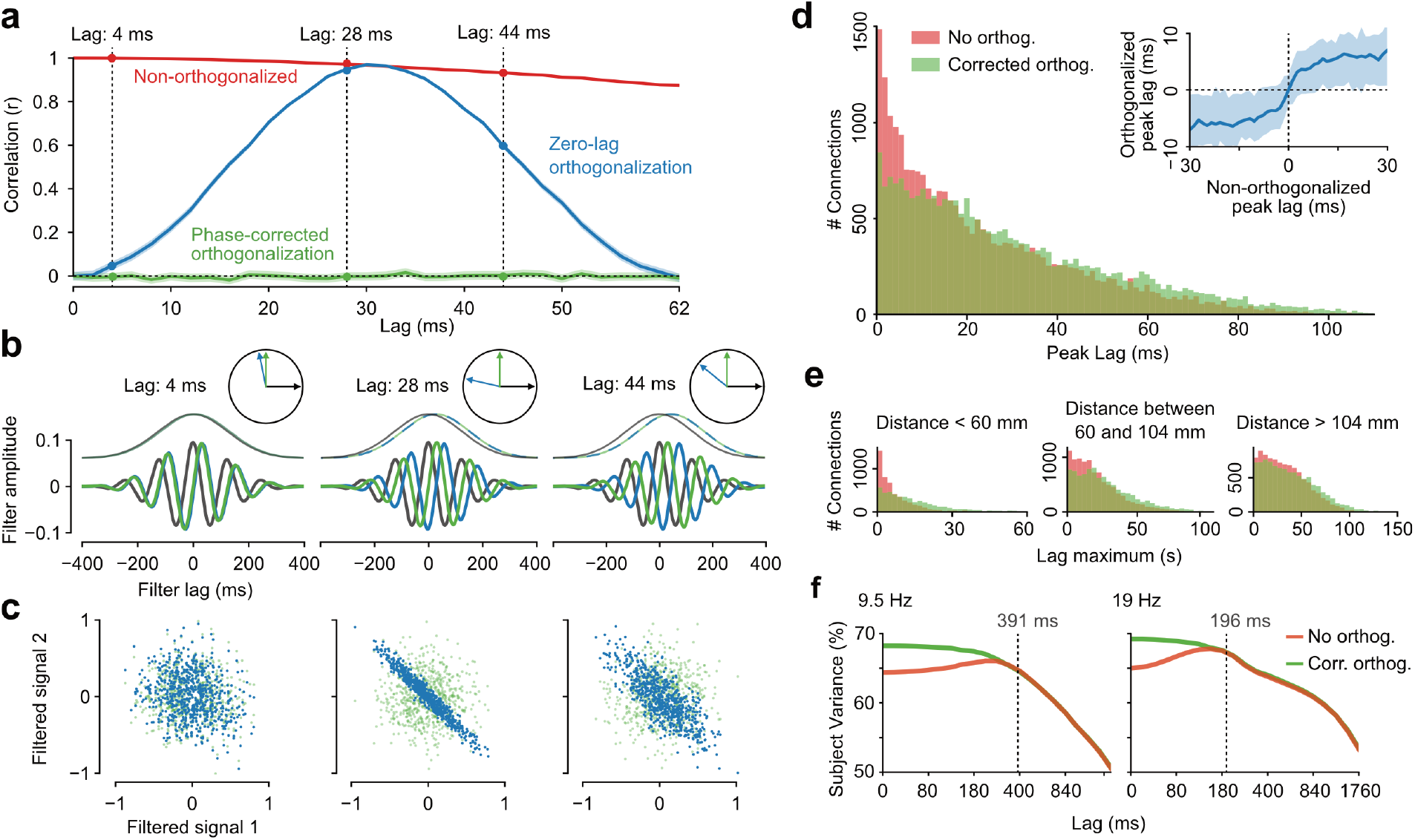
Lag-specific orthogonalization. **a** Orthogonalized and non-orthogonalized correlation of two simulated white noise signals with instantaneous signal mixing. The non-orthogonalized correlation shows a large artifactual correlation (red). Naive zero-lag orthogonalization shows artifactual correlation at non-zero lags (blue). Phase-corrected orthogonalization shows no artifactual correlation (green). **b** Lagged wavelet filters (bottom) and envelopes (top) with (green) and without (blue) orthogonalization phase shift. Reference signal in gray. The insets show the angles between signals. The naively orthogonalized filter rotates and is correlated, depending on lag. The phase-corrected orthogonalization ensures that filters are always orthogonal. **c** Naive zero-lag orthogonalization shows artifactual correlation at non-zero lags (blue). Phase-corrected orthogonalization shows no artifactual correlation (green). **d** Histogram of peak lags with (green) and without (red) orthogonalization in source-reconstructed MEG data at 8 Hz. Inset: Orthogonalization corrects near-zero lags. **e** Stratification of the histogram of peak lags by cortical distance. **f** Comparison of subject-specific variance and noise variance by lag with (green) and without (red) orthogonalization. The dashed lines indicate the temporal width of wavelets. All error-bars denote SEM.

To address this limitation, we introduced a frequency- and lag-dependent phase precession factor into the orthogonalization in the complex wavelet domain (see Methods). This phase precession keeps the relevant wavelet filters orthogonal at the tested lag and eliminated artifactual correlations across the full cross-correlogram (Fig. 1a–c, green). Thus, lag-specific orthogonalization extends zero-lag orthogonalization to amplitude-envelope correlations at non-zero lag.

We next applied the method to resting-state MEG from 89 subjects of the Human Connectome Project ^20^. We performed source reconstruction of cortical activity using beamforming ^21^ and estimated peak lags, i.e. the lags at which amplitude envelope correlation was maximal, for all cortical pairs at 8 Hz (Fig. 1d). Without orthogonalization, peak lags accumulated strongly near zero lag (Fig. 1d, red). This accumulation disappeared after lag-specific orthogonalization, confirming that it was spurious (Fig. 1d, green). The effect was spatially graded: source pairs separated by less than 10 cm were most affected, consistent with the known spatial scale of volume conduction ^18^, whereas pairs separated by more than 10 cm were largely unaffected (Fig. 1e).

Orthogonalization also increased the subject-specificity of short-lag cross-correlations (Fig. 1f). We partitioned crosscorrelation variance into inter-subject, i.e. subject-specific, and inter-session, i.e. noise, variance. For lags longer than the frequency-dependent width of the wavelet (Fig. 1f, dashed lines), this partitioning was nearly identical with and without orthogonalization. In contrast, for lags shorter than the wavelet width, subject-specific variance was higher after orthogonalization. This indicates that artifactual short-lag correlations are less subject-specific than genuine neural coupling.

### Frequency-specific lag threads organize resting-state cortical activity

Having established that lag-specific orthogonalization suppresses spurious correlations while preserving genuine neural signal, we asked whether spontaneous cortical activity follows a structured temporal order – and, if so, whether that order mirrors the brain’s known spatial hierarchy. After removing artifactual correlations, we computed pairwise orthogonalized amplitude envelope cross-correlograms across all cortical sources and 25 frequency bands from 2 to 128 Hz. For each source pair, we extracted the peak lag, yielding an antisymmetric time-delay matrix for each frequency band (Fig. 2a).

**Fig 2.**
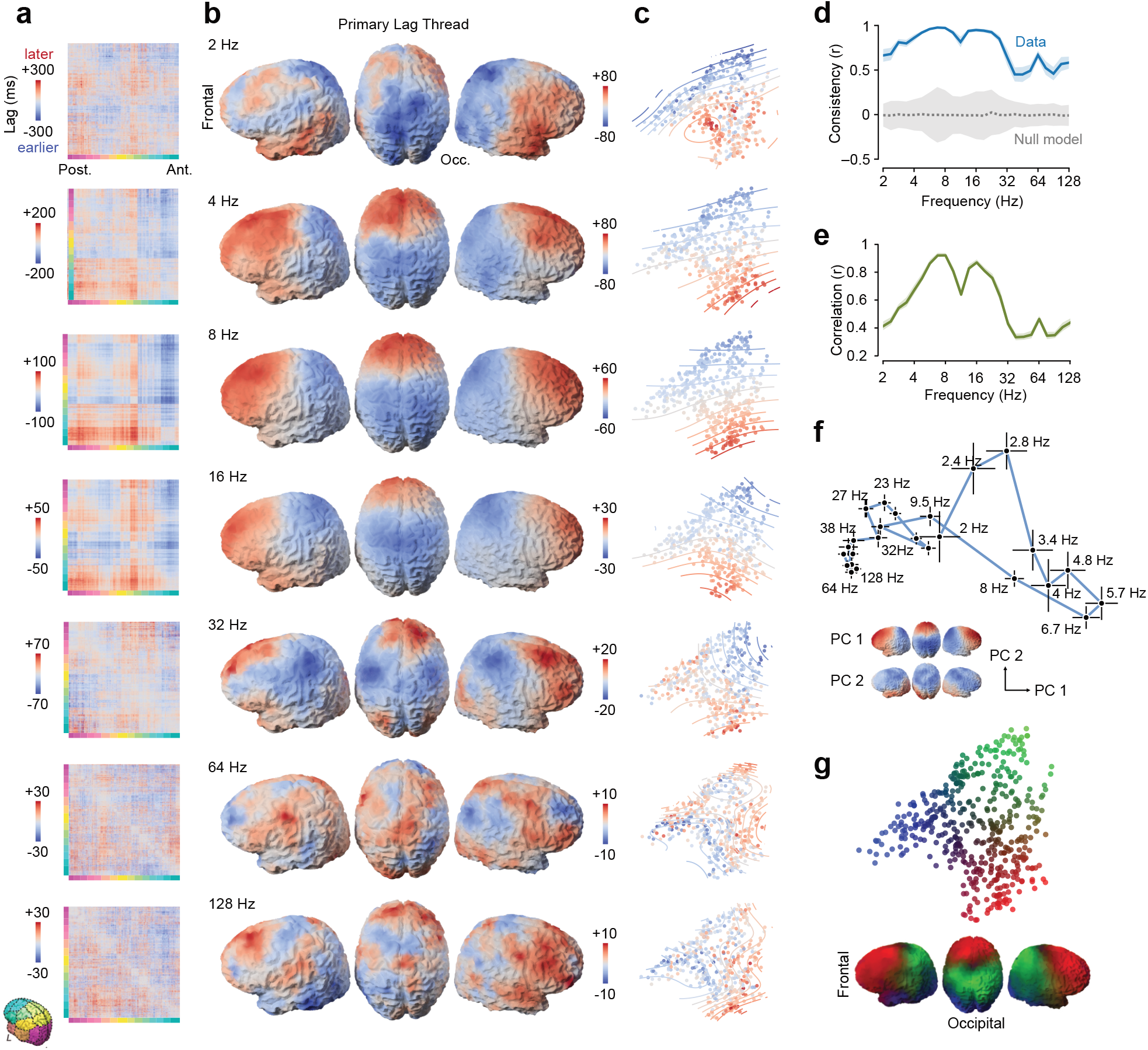
Frequency-specific lag threads. **a** Matrix of peak cross-correlation lags across pairs of cortical regions. Regions are sorted from posterior to anterior, as indicated in the inset. **b** Primary lag thread for each frequency range defined as the average lag of each region to all other regions. **c** Primary lag threads of all cortical regions embedded in the tripolar gradient shown in (g). Curved lines show level sets of a Gaussian kernel estimate across the two-dimensional embedding space. **d** Inter-subject consistency of lag threads quantified as the correlation between bootstrap samples across subjects. The gray line shows a subject-level null model (sign flip). **e** Correlation between the peak-lag matrix (a) and its rank-one approximation. **f** Two-dimensional embedding (PCA) of lag threads across frequencies. **g** Two-dimensional embedding (PCA) of lag threads across cortical regions (colored dots). Cortical locations are RGB color coded by barycentric coordinates of a triangular convex hull of embedded points as indicated on the cortical surface. All errorbars denote SEM across subjects.

For each frequency, we derived the primary lag thread, i.e. the dominant spatial map of temporal ordering, by averaging each row of this matrix. This is equivalent to a least-squares rank-one approximation (Fig. 2b). This rank-one structure is the basis for interpreting lag threads as a temporal hierarchy. If one region tends to precede a second region, and the second tends to precede a third, the full time-delay matrix can be approximated by the differences between regional positions along a single temporal axis. The spatial structure of primary lag threads was bilaterally symmetric and consistent across subjects for all frequencies. Correlations of lag threads across bootstrap samples of the 89 subjects exceeded correlations expected by chance (Fig. 2d; all frequencies p < 0.01, FDR corrected; permutation test). Activity was consistently earliest in occipital and parietal cortex and latest in frontal and temporal cortex, following a progression from unimodal sensory regions toward transmodal association cortex that broadly mirrors the principal spatial gradient of the brain ^1,2^ (Fig. 2b).

The amount of pairwise lag variance captured by the primary lag thread depended strongly on frequency (Fig. 2e). In the alpha (around 8 Hz) and beta (around 16 Hz) bands, the primary thread explained up to 85% of lag variance, indicating a highly structured, near-transitive temporal hierarchy. At the edges of the investigated frequency range, explained variance decreased to approximately 11%.

Principal component analysis of lag threads across frequencies (Fig. 2f) revealed a smooth frequency dependence. Across frequencies, lag patterns spanned a multi-dimensional space, with distinct temporal precedence hierarchies at different frequencies.

To characterize the spatial variation of temporal precedence across the cortex, we performed a principal component analysis of the primary lag threads across cortical sources. This projection revealed a tripolar structure of frequency-dependent lags with poles in occipitotemporal, sensorimotor, and frontal cortex (Fig. 2g). This tripolar structure is highly similar to embeddings derived from structural data, fMRI functional connectivity, and other dynamical and structural measures ^2,22–24^. Since the dimensions in our embedding are relative time delays (Fig. 2c), the MEG-based discovery of a similar spatial patterns adds a temporal dimension to the gradient architecture of human cortical organization ^25–28^.

### Lag threads covary with handedness and age

Although lag threads were largely bilaterally symmetric (Fig. 2b), they also showed a consistent lateral asymmetry in the resting state: Averaged across all frequencies, right hemisphere activity tended to precede activity in the left hemisphere by 8 ms (± 1.5 ms). This effect was strongest around 11 Hz and statistically significant (p < 0.004, FDR-corrected; see also Fig. 6b).

If lag threads reflect neurobiological organisation rather than an analytic artefact, they should be sensitive to individual neurobiological differences whose cortical signatures are well established, such as the hemispheric lateralization of handedness and the progressive cortical changes accompanying ageing. We tested whether the observed inter-hemispheric asymmetry reflects handedness by correlating the inter-hemispheric component of the lag thread with Edinburgh Handedness Inventory (EHI) scores ^29^. Inter-hemispheric lag asymmetry was significantly correlated with average handedness (Fig. 3a; p < 0.05 FDR corrected), maximally explaining 18.8% of lag thread variance across bootstrap samples at 16 Hz. At this frequency, more right-handed subjects showed reduced lag thread asymmetry in leave-one-out pseudovalues (Fig. 3b; r = 0.22; p = 0.036). The correlation was strongest in occipital and parietal cortex (Fig. 3c). Thus, MEG lag thread asymmetry is sensitive to handedness, although the anatomical distribution suggests that it may reflect visuospatial or attentional lateralization rather than motor lateralization alone.

**Fig 3.**
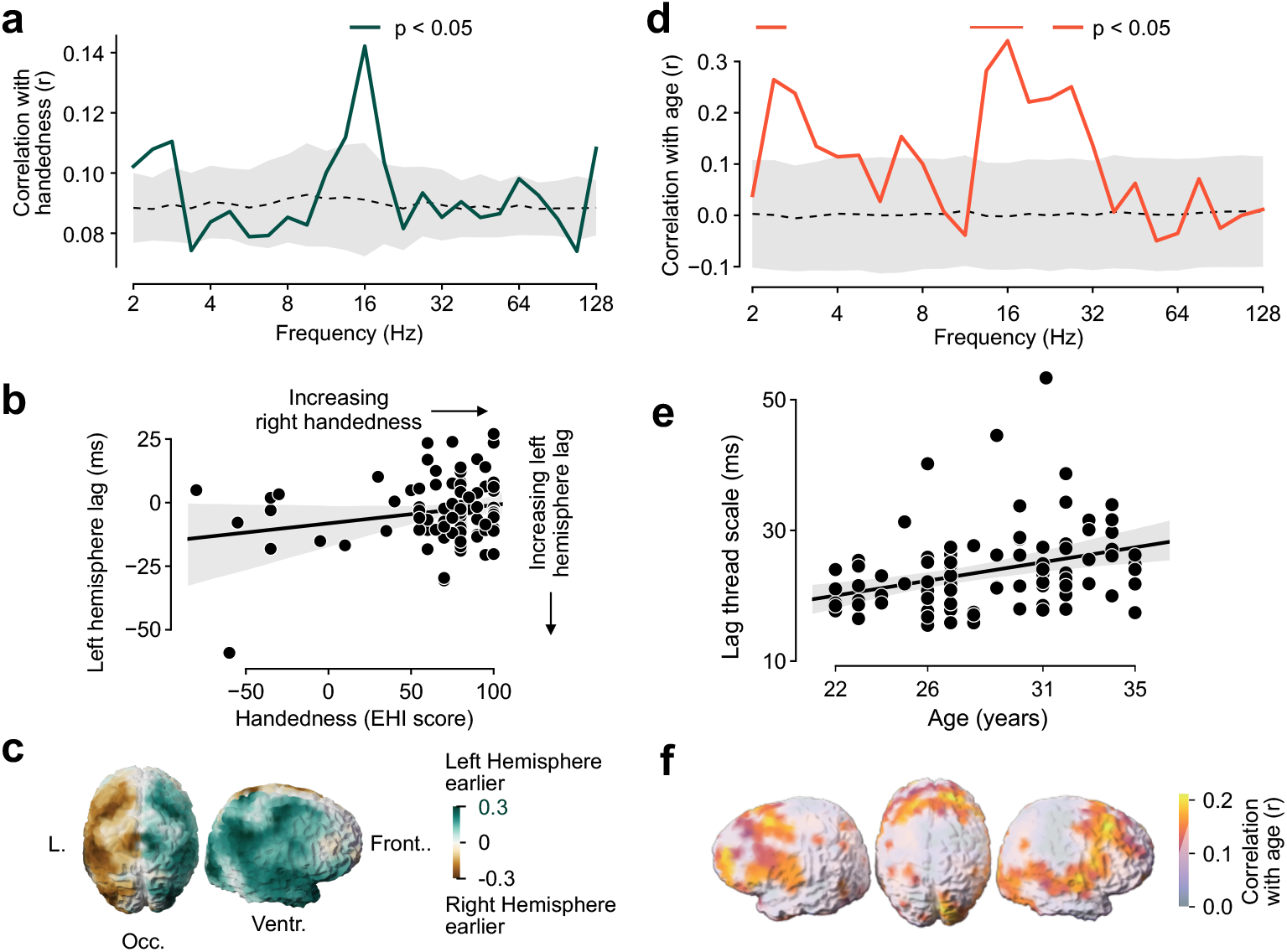
Lag threads are modulated by subject handedness and age. **a** Correlation between average EHI handedness score of subjects and the average inter-hemisphere lag across bootstrap samples. Shaded gray area shows the standard deviation of correlations in a subject-level permutation null model. **b** Relation between subject handedness and the average left-hemisphere lag at 16 Hz. **c** Cortical distribution of the correlation between handedness and hemispheric asymmetry. **d** Correlation between the temporal scale (standard deviation) of the primary lag thread and subject age. Shaded gray area shows the standard deviation of correlations in a subject-level permutation null model. **e** Relation between subject age and the temporal scale (standard deviation) of the primary lag thread at 16 Hz. **f** Cortical distribution of the correlation between subject age and the temporal scale of the primary lag thread at 16 Hz. Non-masked sources are significant at p < 0.05 (FDR corrected) with respect to a subject-level permutation null model.

We next asked whether lag threads were related to age. The overall temporal scale of the primary lag thread, quantified as the root mean square of the lag thread across the brain, was significantly correlated with subject age around 2.4 Hz and from about 11 Hz to 27 Hz (Fig. 3d; p < 0.05 FDR corrected; permutation test). The correlation was strongest at 16 Hz (r = 0.34; p = 0.001) and positive, indicating that the scale of temporal lags grows within this young-adult sample (Fig. 3e). The cortical sources contributing most to this effect were located bilaterally in frontoparietal and temporal cortex (Fig. 3f).

### MEG and fMRI lag threads share subject-specific spatial variance

If fMRI lag threads have a neural basis, then electrophysiological and hemodynamic lag maps should not only share spatial structure at the group level, but should covary as a subject-specific signature. We tested this directly in the same 89 individuals by extracting primary lag threads from their resting-state BOLD data. We resampled the cortical fMRI time series to the MEG 457-source model using a 5 mm Gaussian kernel and computed cross-correlograms at lags up to 20 TRs (14.4 s) (Fig. 4a). The fMRI primary lag thread recapitulated known patterns, with maximum average lags of approximately 0.5 s. These lags were substantially longer than MEG envelope lags, consistent with the different time-scales of the two signals.

**Fig 4.**
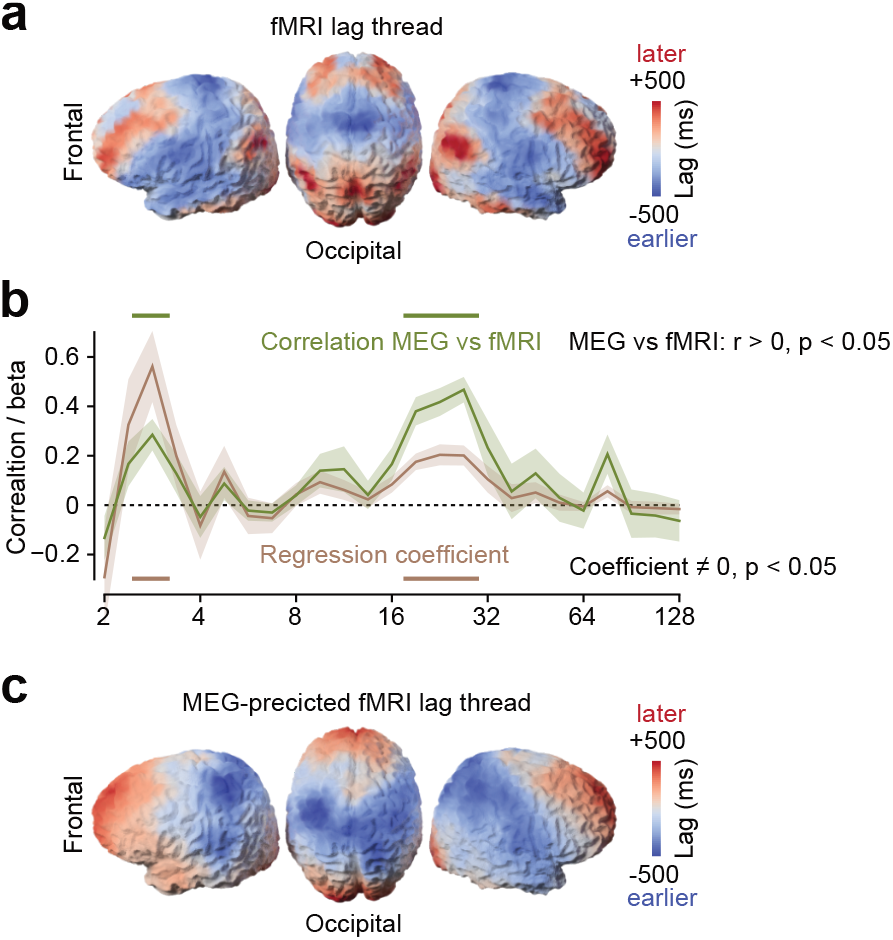
MEG and fMRI lag threads are correlated. **a** Primary lag thread of the HCP resting state fMRI data, resampled to the MEG source model. **b** Spatial correlation between the fMRI and MEG lag threads by frequency (green curve). The brown curve shows the coefficients of a ridge regression (λ = 10^3^) to predict fMRI lag threads from a linear combination of MEG lag threads in all 25 frequencies. Error-bars denote SEM across subjects. **c** Average MEG-based prediction of fMRI lag thread.

Single-frequency MEG lag threads were spatially correlated with the fMRI lag thread across several frequencies (Fig. 4b; p < 0.05, FDR corrected), with a maximum Pearson correlation of r = 0.47 at 26.9 Hz. This spatial correspondence captures the similarity between group-level MEG and fMRI lag maps and is therefore distinct from subject-specific covariance. A regularized multi-frequency linear model (ridge regression, λ = 10^3^) predicting the fMRI lag thread from all 25 MEG frequency bands explained 21.2% of fMRI lag spatial variance. Regression coefficients were significantly different from zero at several frequencies (Fig. 4b; p < 0.05, FDR corrected) and weighted most strongly toward low MEG frequencies, in particular around 2.8 Hz. The difference between the peak single-frequency correlation and the low-frequency ridge weights suggests that spatial similarity and multivariate prediction capture partially distinct aspects of the MEG-fMRI relationship.

Critically, the cross-modal correspondence also contained subject-specific covariance. Predictive fit was significantly higher when MEG and fMRI lag threads were derived from bootstrap samples containing the same subjects than from independently resampled subject sets (p < 0.001, permutation test). This matched-sample advantage was small in absolute terms (∼0.8% improvement in variance explained), consistent with the modest contribution of any single individual to a group bootstrap average, but it was highly consistent across permutation samples, indicating that individual variation in MEG amplitude lag structure and fMRI propagation sequences is correlated beyond the shared group-level spatial pattern. This convergence supports a neural contribution to fMRI lag structure.

### Lag threads are selectively reconfigured during cognitive task performance

Lag threads could be fixed by anatomy or dynamically adapt to cognitive demands. We found evidence for both. We compared resting-state lag threads with lag threads during active task performance in two complementary datasets. In the HCP working-memory dataset ^20^ (Fig. 5; visual n-back task with right-hand motor responses; N = 89 subjects), the overall lag thread structure was largely preserved (Fig. 5). Resting-state and working-memory primary lag threads were significantly correlated across a broad frequency range (Fig. 5a, green, p < 0.05, FDR corrected) and reached a peak correlation of r = 0.93 around 8 Hz.

**Fig 5.**
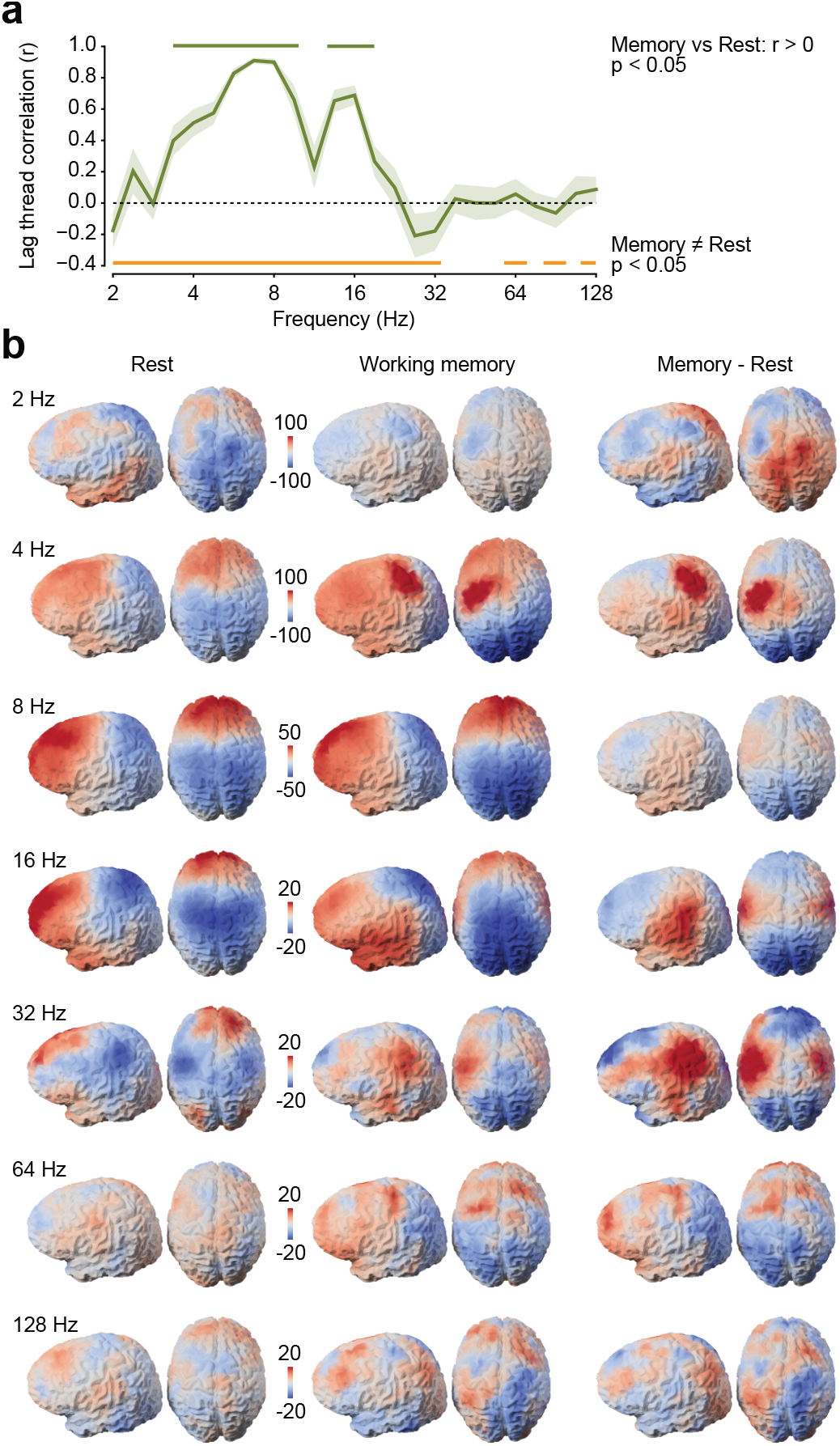
Lag threads during working memory. **a** Correlation of primary lag threads between resting state and working memory. Shaded region denotes SEM across subjects. The green bar indicates frequencies with significant correlation (p < 0.05, FDR corrected, t-test). The orange bar indicates frequencies with a significant difference of lag threads between rest and working memory (p < 0.05, FDR corrected, permutation test). **b** Primary lag threads during rest, working memory and their difference.

Nevertheless, we observed selective changes with task at several frequencies (Fig. 5a, orange, p < 0.05, FDR corrected; permutation test). At 4 Hz, left sensorimotor cortex shifted to a later position in the temporal sequence, consistent with preparation and execution of right-hand button presses, whereas primary visual cortex shifted toward earlier temporal precedence, consistent with enhanced sensory drive (Fig. 5b, right column). Similarly, at 32 Hz, sensorimotor cortex shifted later bilaterally and frontoparietal regions shifted earlier. Thus, the task induced targeted reconfiguration of the temporal hierarchy on top of a largely preserved resting-state scaffold.

The n-back task involved both sensory stimulation and motor responses, which could independently drive the observed lag thread changes. Could reconfiguration occur during purely internal cognition, where neither confound is present? To address this, we analyzed a second dataset from 28 subjects who performed a covert autobiographical memory recall task while fixating (Fig. 6). Subjects were instructed to silently recall the entire day from the moment they woke up until the start of the recording, revisiting as many details as possible in chronological order. Crucially, this condition was matched to resting state in terms of sensory input and motor output, isolating any effect of internally generated cognitive engagement.

**Fig 6.**
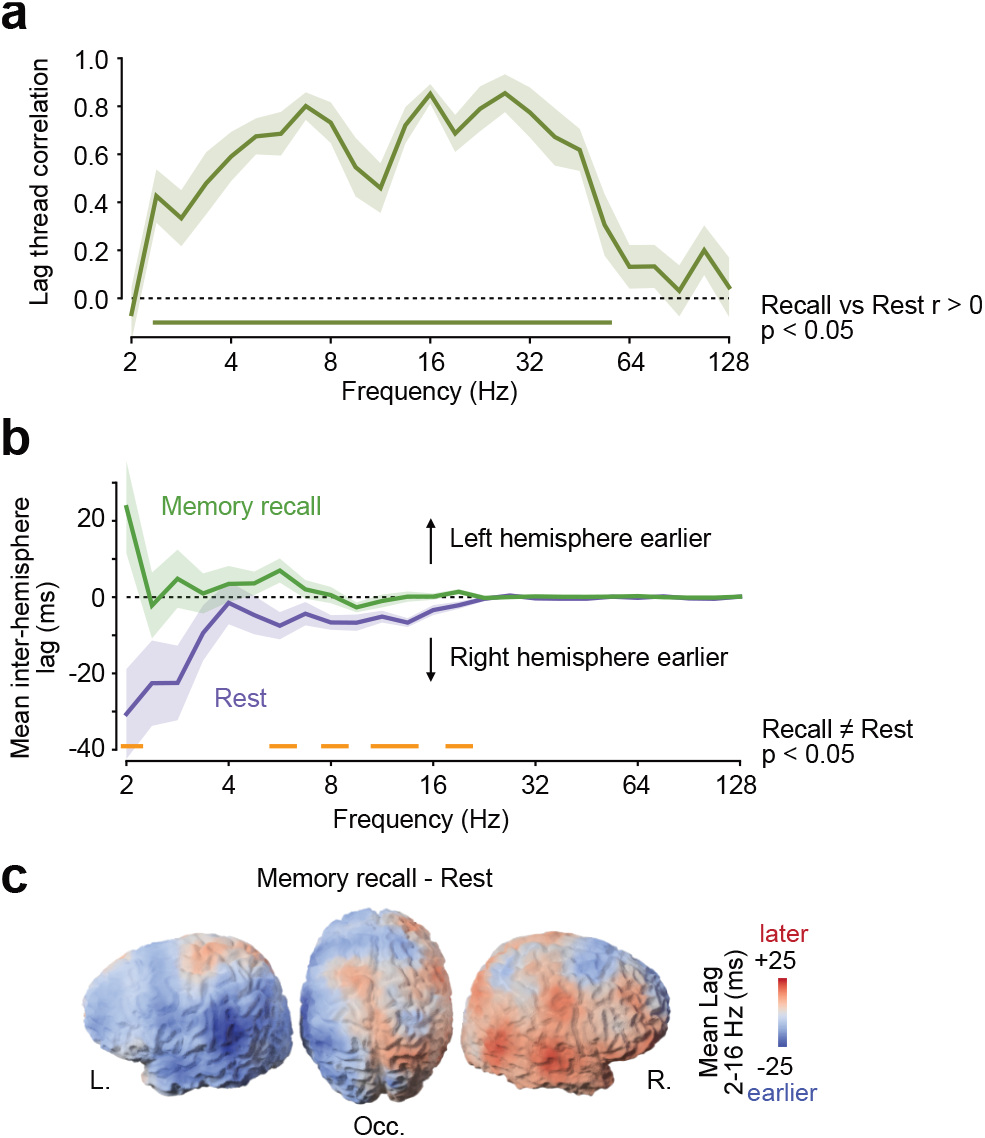
Lag threads during covert memory recall. **a** Correlation of primary lag threads between resting state and covert memory recall. The shaded area denotes SEM across subjects. The green bar indicates frequencies with significant correlation (p < 0.05, FDR corrected, t-test). **b** Average lag between the left and right hemisphere as a function of frequency during rest and memory recall. The orange bar indicates frequencies with significant difference between conditions (p < 0.05, FDR corrected, t-test). **c** Cortical distribution of the difference of lag threads between memory recall and rest averaged from 2 to 16 Hz.

The overall correlation of lag threads between rest and task was lower in this dataset than in the HCP comparison, consistent with the smaller sample and the more variable cognitive state during uncued recall (Fig. 6a). Nevertheless, correlations between conditions were significantly positive across a broad frequency range (Fig. 6a, p < 0.05 FDR corrected).

Directly contrasting rest and memory recall across all frequencies did not yield a significant effect (p > 0.05, FDR corrected). However, memory-recall had a prominent effect on the hemisphere-lateralization of lag threads. Memory recall induced a reduction, and at several frequencies between 2 Hz and 19 Hz even a reversal, of the resting-state right-hemisphere temporal precedence (Fig. 6b, p < 0.05, FDR corrected). The spatial pattern of this difference (Fig. 6c) involved regions consistent with the left-lateralized language network, suggesting that engagement of language-related processes during memory recall may shift the temporal hierarchy toward left-hemisphere precedence. Together, these results show that lag thread reconfiguration extends beyond externally driven sensory-motor tasks to internally generated cognition.

## Discussion

We identify frequency-specific electrophysiological lag threads as a temporal hierarchy of the human cortex. In resting-state MEG, cortical amplitude fluctuations followed a reproducible temporal ordering that was most coherent in the alpha and beta bands and broadly aligned with the sensory-to-association organization of the cortex ^1,2^. This near-transitive lag structure shows that pairwise delays are not distributed randomly across the connectome but can be summarized by a dominant cortical thread from earlier to later regions. Lag thread maps thereby add a temporal dimension to macroscale cortical gradients: they describe the direction and delay of activity fluctuations across cortex.

This result was enabled by lag-specific orthogonalization. Conventional pairwise orthogonalization ^18^ removes instantaneous leakage effects, but does not guarantee orthogonality after a temporal shift and therefore leaves oscillatory artifacts in non-zero-lag envelope correlations. By introducing a frequency- and lag-dependent phase precession in the complex wavelet domain, the present approach restores orthogonality at each tested lag. It thereby makes lagged amplitude-envelope correlations accessible in source-reconstructed MEG, precisely in the lag range in which volume conduction and signal leakage are most problematic. The method extends established leakage-correction approaches from static envelope coupling to temporally ordered envelope dynamics ^18,19^.

The correspondence between MEG and fMRI lag threads provides a link between fast electrophysiological dynamics and slower hemodynamic propagation sequences. fMRI lag threads have been proposed to organize intrinsic activity, but their interpretation is limited by the indirect nature of the BOLD signal and by possible regional variation in neurovascular coupling ^8–10^. The finding that MEG and fMRI lag maps share spatial structure, and that this correspondence contains subject-specific covariance, supports a neural, instead of vascular, origin of fMRI lag structure. The strong weighting of the cross-modal model toward very low MEG frequencies (approximately 2.8 Hz) is consistent with the known coupling between infraslow neural fluctuations and the BOLD signal ^30,31^. At the same time, the correspondence was partial. MEG and fMRI operate on different timescales, were recorded non-simultaneously, and likely capture overlapping but non-identical aspects of intrinsic dynamics. Thus, the cross-modal result provides evidence for a shared component of multiscale temporal organization, not for a one-to-one mapping between electrophysiological and hemodynamic lags.

Several interpretational limits are important. Amplitude-envelope lags describe the temporal ordering of power fluctuations, not the direction of synaptic transmission or information transfer. Thus, posterior-to-anterior lag structure should not automatically be interpreted as directed information flow. Similar temporal ordering could arise if cortical regions respond with different latencies to shared network fluctuations, for example because of regional differences in excitability, intrinsic timescales, structural connectivity, or neuromodulatory influences. Likewise, lag threads are related to, but distinct from, cortical traveling waves ^32–35^. Traveling waves describe cycle-by-cycle phase gradients, whereas lag threads describe the ordering of temporal amplitude envelopes ^36,37^ (i.e., phase velocity vs. group velocity ^38^). Propagating oscillatory bursts or time-irreversible state transitions and dynamics ^12,39–41^ could contribute to both phenomena, but envelope lag structure can also arise without a continuous phase-defined wave. Thus, the present measure provides a complementary description of large-scale temporal organization rather than direct evidence for directed signaling or traveling-wave propagation. Regardless of the mechanism, the time-directed nature of the non-zero lags is a marker of time-irreversible, non-equilibrium cortical dynamics ^42,43^. Unlike a system in thermal equilibrium whose dynamics are symmetric in time, the cortex sustains a directed flow of activity that cannot arise from passive fluctuations alone – a property shared with, but not reducible to, either traveling waves or directed synaptic drive.

The individual difference and task results show that lag threads are not fixed anatomical constraints. Resting-state inter-hemispheric lag asymmetry covaried with handedness, suggesting sensitivity to established lateralization traits. The age-related increase in the temporal scale of lag threads is also intriguing, but the HCP young-adult sample covers only a narrow age range. Broader developmental and ageing cohorts will be required to determine whether this effect reflects a true lifespan trajectory.^3^ The task analyses provide the clearest evidence for functional reconfiguration. During visual working memory, the canonical hierarchy was largely preserved but selectively shifted in regions expected from sensory input and right-hand responses. During covert autobiographical memory recall, changes in inter-hemispheric precedence were consistent with engagement of left-lateralized language-related networks. Together, these findings suggest that lag threads combine a stable cortical scaffold with state-dependent modulation.

This work also has implications for clinical translation. Previous fMRI studies have shown that lag structure is focally disrupted in autism spectrum disorder, other neuropsychiatric conditions ^4,7,44^, and pharmacological interventions ^45,46^. MEG-based lag analysis could provide a complementary electrophysiological window into these disruptions. It is applicable where fMRI is contraindicated or poorly tolerated, and – crucially – it affords time-resolved discrimination between neural timing disruptions and potential neurovascular confounds. Furthermore, frequency-specific lag maps may reveal whether disruptions are confined to particular oscillatory bands, adding a dimension not accessible to fMRI. The present results provide the methodological basis and cross-modal validation for this approach. However, individual-level reliability remains a key challenge before clinical applications are feasible.

In summary, lag-specific orthogonalization reveals a previously obscured temporal axis of human cortical organization. Frequency-specific MEG lag threads are reproducible across individuals, partially shared with fMRI propagation structure, sensitive to individual traits, and reconfigured by cognitive state. These properties suggest that the temporal ordering of cortical amplitude fluctuations is a genuine and functionally relevant feature of large-scale human brain dynamics.

## Acknowledgements

We thank Joerg Hipp and Anna-Antonia Pape for recording part of the Tübingen (memory recall) MEG dataset. We thank Markus Siems for help with preprocessing HCP data. This study was supported by the European Research Council (ERC; https://erc.europa.eu/) CoG 864491 (M.S.) and by the German Research Foundation (DFG; https://www.dfg.de/) project SI 1332/6-1 (SPP 2041) (M.S.).

## Author contributions

P.H.: Conceptualization, Methodology, Software, Formal analysis, Visualization, Writing – original draft, Writing – Review & Editing. J.G.: Investigation, Data curation. Writing – Review & Editing. M.S.: Conceptualization, Supervision, Resources, Project administration, Funding acquisition, Writing – original draft, Writing – Review & Editing.

## Data and code availability

The HCP MEG and fMRI data used in this study are publicly available through the Human Connectome Project at https://db.humanconnectome.org (S900 release).

## Materials and Methods

### Orthogonalization of lagged amplitude envelope correlations

Our method for computing cross-correlations with pairwise, time- and frequency specific orthogonalization is based on a well-known method for the zero-lag case ^18^. Given two signals *x*_1_(*t*) and *x*_2_(*t*) and their complex-valued time-frequency transforms *y*_1,*f*_(*t*) and *y*_2,*f*_(*t*), where *f >* 0 is the base frequency^1^, a version of the zero-lag orthogonalization method consists in computing the correlation between

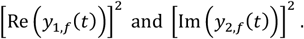

This is effective in eliminating artifactual correlation at lag zero but not at non-zero lags (Fig. 1a-c). Note that the original method instead computes 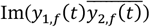 which has a similar effect.

The zero-lag orthogonalization method is effective because the signals

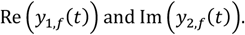

are, in the case of a Morlet-based continuous wavelet transform, the convolutions of the original signals *x*_1_(*t*) and *x*_2_(*t*) with the real and imaginary parts of the Morlet wavelet *w*(*t*), respectively (because *x*_1_(*t*) and *x*_2_(*t*) are real-valued). Because the real-valued filters

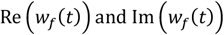

are orthogonal in the standard Euclidean inner product

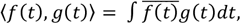

any zero-lag mixing between *x*_1_(*t*) and *x*_2_(*t*) acts independently on the filtered signals Re(*y*_1,*f*_ (*t*)) and Im(*y*_1,*f*_(*t*)). At non-zero lags, the method is only effective when the cross-correlation between Re(*w*_*f*_(*t*)) and Im(*w*_*f*_ (*t*)) is zero. However, this is only the case for lags that are integer multiples of the inverse base frequency 1*/f*.

To correct for signal mixing at non-zero lags, we must introduce a phase factor *ϕ*(*f,l*) depending on the lag *l* ∈ ℝ and the base frequency *f*. The phase factor *ϕ*(*f,l*) is chosen such that the cross-correlation at lag *l* of the real part of *w*_*f*_ (*t*) and the imaginary part of the time-shifted and phase-multiplied wavelet *ϕ*(*f, l*) · *w*_*f*_ (*t* + *l*) is zero:

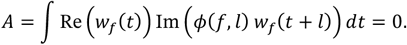

The fact that such a phase factor *ϕ*(*f, l*) exists for all frequencies *f* > 0 and lags *l* ∈ ℝ follows from the intermediate value theorem as follows:

From the formula above, it is clear that if we set the phase factor *ϕ*′ (*f, l*) = −*ϕ*(*f, l*), we obtain *A*′ = −*A*. By the intermediate value theorem on the circle, there has to be some value *ϕ* of norm 1 for which A = 0.

In case that *w*_*f*_(*t*) is a complex Morlet wavelet (Gabor filter), we can derive a simple formula for the phase. Define a Gaussian envelope as

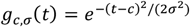

and a general real-valued Gabor filter as

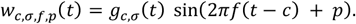

An elementary computation yields the following formula for the product of two wavelet filters:

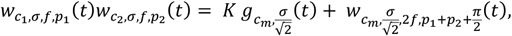

where 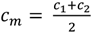 and

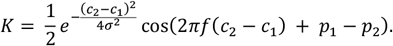

For the two wavelet filters to be orthogonal, the integral of 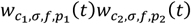 over the real line must be zero. Now 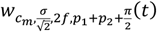 always integrates to zero since it is antisymmetric around *c*_*m*_, and the Gaussian 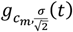 always has a positive integral, so the cosine term in K must vanish for the wavelets to be orthogonal. This is the case only if 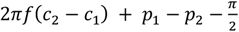 is a multiple of *π*. Since we have *p*_1_ = *c*_1_ = 0, we can achieve this by simply setting 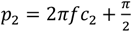. In terms of the phase factor *ϕ*(*f, l*), this means that we can set

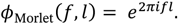

We used this simple formula to orthogonalize Morlet wavelet amplitudes in the following.

Including the phase factor, we may then compute the cross-covariance by

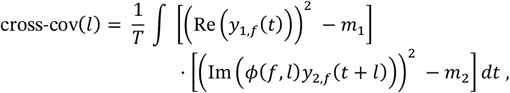

where *m*_1_ and *m*_2_ denote the time average of (*Re*(*y*_1_,_*f*_(*t*)))^2^ and (*Im*(*ϕ*(*f,l*)y_2_,*f*(*t* +*l*)))^2^, respectively. The cross-correlation is obtained by dividing each time series by its standard deviation. Setting *ϕ*(*f*,0) = 1 recovers the original orthogonalization method ^18^ for lag zero.

We introduce one additional improvement for the computation of amplitude envelope cross-correlations, which concerns the spatial content of the source-reconstructed M/EEG signal. The result of beamforming is often a vector-valued, three-dimensional current or dipole moment 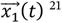.Conventionally, this signal is projected onto the first principal component of its spatial time course (that is, one sets 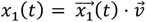 where 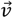 is one of the unit vectors maximizing the Frobenius norm 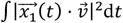. However, this is arbitrary and discards data. Instead, we here compute the wavelet transforms 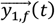 vectorially (that is, independently on each component of the vector). We then compute

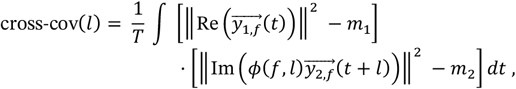

where ‖·‖ denotes the Euclidean norm in three dimensions, and *m*_1_ and *m*_2_ denote the time average to ‖*Re*(*y*_1_,_*f*_(*t*))‖^2^ and ‖*Im*(*ϕ*(*f,l*)y_2_,*f*(*t* +*l*))‖^2^ respectively. This approach improved the SNR of the cross-correlograms.

### Testing the orthogonalization method on simulated data

To test the orthogonalization method on simulated data, we generated two independent white noise signals *x*_1_(*t*) and *x*_2_(*t*) with a sampling rate of 1000 Hz, representing uncoupled neural data. Artifactual mixing of strength *λ* ∈ [0,1] was simulated by setting *y*_1_(*t*) = (1 *-λ*)*x*_1_(*t*)+ *λx*_2_(*t*) and *y*_2_(*t*) = *λx*_1_(*t*) + (1 *-λ*)*x*_2_(*t*). For a frequency of *f* = 8 Hz, we computed the complex continuous wavelet transform (CWT) with a Morlet wavelet with a full width at half maximum (FWHM) of 465 ms for the signals *y*_1_(*t*) and *y*_2_(*t*).

Since the underlying signals *x*_1_(*t*) and *x*_2_(*t*) were uncorrelated white noise signals, the expected ground truth cross-correlation of our simulation was zero, and any observed cross-correlation is an artifact of signal mixing. We computed the amplitude envelope cross-correlation between *y*_1_(*t*) and *y*_2_(*t*) using three algorithms: 1) Cross-correlation without orthogonalization of the absolute values of the CWTs of *y*_1_(*t*) and *y*_2_(*t*). 2) Naive orthogonalization using the lag zero method. We correlated the absolute value of the real part of the CWT of *y*_1_(*t*) with that of the imaginary part of the CWT of *y*_2_(*t*), and vice versa. At lag zero, this method is effective at eliminating zero-lag artifactual coupling ^18^. 3) Corrected cross-correlation using a lag-specific phase offset, as described in the previous section.

We computed the cross-correlations according to all three methods for 104 independent generations of the underlying signals *x*_1_(*t*) and *x*_2_(*t*).

### MEG data and preprocessing

We used MEG data from two datasets. For most of the analyses (Fig. 1–4), we used publicly available MEG data from the Human Connectome Project (HCP) S900 release ^20^. We only analyzed data from those *N* = 89 subjects (41 female, aged 22–35 years) where data from all three MEG resting state sessions were available. We also used MEG recordings from the working memory task, in which subjects performed a visually presented n-back task of varying difficulty, reporting their results via a right-hand button press. HCP data were recorded using a whole-head Magnes 3600 scanner (4D Neuroimaging, San Diego, CA, USA) in a magnetically shielded room (22). Structural, T1-weighted MRI scans were acquired using a Siemens 3T Skyra scanner, with a spatial resolution of 0.7 mm.

In addition, for the results presented in Figure 6, we analyzed data recorded at the MEG Center Tübingen, in which subjects performed a covert autobiographical memory recall task without overt sensory cueing or motor responses. This dataset comprised N = 28 healthy subjects (17 female, mean age 26.5 years). For each subject, 10 minutes of resting-state MEG and 10 minutes of task MEG were recorded with 275 channels at a sampling rate of 2,343.75 Hz (Omega 2000, CTF Systems, Inc., Port Coquitlam, Canada). In both conditions, participants were seated upright in a dimly lit magnetically shielded chamber and fixated a central fixation point. In the memory recall condition, subjects were instructed before the recording to silently recall the entire day from the moment they woke up until the start of the experiment, revisiting as many details as possible in chronological order. The recordings were approved by the local ethics committee and conducted in accordance with the Declaration of Helsinki. All participants gave written informed consent before participating.

For the HCP data, we used preprocessed MEG data provided by the HCP project ^39^. This preprocessing pipeline includes the removal of noisy or bad channels, as well as the removal of segments of bad data and non-brain physiological artifacts using independent component analysis (ICA) ^39,40^. A similar process was used for the locally recorded dataset.

For both datasets, we followed 31 for physical forward modeling of signals. MEG sensors were aligned to individual anatomy in FieldTrip ^41^. The forward model was based on a single-shell head model based on individual T1-weighted structural MRI scans ^42^.

We modeled the source of the measured magnetic fields and field gradients by a superposition of three-dimensional current elements at 457 locations in the brain. The source locations were spaced equally at a distance of around 1.2 cm at approximately 0.7 cm below the cortical surface ^43^. The forward model of magnetic fields generated by the source currents was computed using a boundary element method using a single-shell head model based on the anatomical MRI scan ^42^. We reconstructed source currents using an LCMV beamformer ^21^.

We computed a time-frequency representation at 25 frequency bands logarithmically spaced between 2 and 128 Hz (four bands per octave) using a complex-valued Morlet CWT. The time and frequency resolution were frequency dependent, with FWHM-time = 1.9 s and FWHM-frequency = 0.17 Hz for the lowest band (2 Hz) and FWHM-time = 29 ms and FWHM-freq = 21.9 Hz for the highest band (128 Hz). We computed the wavelet transform with a sampling rate of 50 Hz.

### Computation of cross-correlograms and estimation of peak lags

We computed pairwise orthogonalized cross-correlations in each frequency band between each pair of sources in our 457-source model. We computed cross-correlograms at lag zero and each of a geometrically increasing sequence of 33 non-zero lags ranging from 20 ms to 18.9 s.

An important quantity for the cross-correlogram is the lag at which the cross-correlation is maximal. Previous studies have estimated this lag in fMRI by fitting a parabola to the three points around the lag which maximizes the cross-correlation ^8^. Here, we proceeded similarly by performing a least-squares fit of a Gaussian 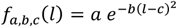 to the central 21 points of our cross-correlograms, the value *c* ∈ ℝ being the lag peak.

While the lag peak and other statistics can be computed easily and reliably on the group average cross-correlation, it is difficult to obtain reliable estimates from the 15 minutes of data available for a single subject. Therefore, in order to obtain statistical measures at the group level, we used a bootstrap approach. For *n*_bt_ = 100 bootstraps, we sampled at random, with replacement, 267 from among the 267 resting state sessions and averaged the cross-correlation from these sessions. We separately performed the Gaussian least-squares fit on each bootstrap average. Thus, the standard deviation of peak lag and other statistics approximates the standard error of these quantities across subjects.

### Transitivity and lag threads

Using the method described in the previous section, we obtain a matrix *A* of peak lags of shape *n*_src_ *× n*_src_, where *n*_src_ = 457 is the number of cortical sources, the time delay (TD) matrix ^8^. Because the cross-correlogram for the pair (*i, j*) is the temporal reversal of the cross-correlogram for (*j,i*), this matrix is antisymmetric (*A*^*T*^ = *–A*).

To test whether the pattern of peak lags is consistent across subjects, we compared the average correlation of lag threads in bootstrap samples (see section “Computation of cross-correlograms and estimation of peak lags”) to a null model describing the hypothesis that no consistent lag thread across subjects exists. For this null model, we randomly changed the time direction of cross-correlations (and thus the sign of the peak lag) with probability 0.5 for each subject for 100 independent bootstrap samples. We then compared the average correlation between bootstrap samples with (null model) and without flipping of time directions (gray and blue lines in Fig. 2d.)

A central finding in fMRI lag threads is that this time delay matrix is often well-described by the difference of two vectors,

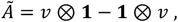

Where *ν* is an n_src_-dimensional vector and **1** is a vector of the same dimension whose entries are all 1. For antisymmetric matrices *A*, we can compute the best fit for such a vector by simply averaging the rows of the matrix ^8,44^. We refer to the vector *v* as the primary lag thread. We computed the correlation between flattened matrices *A* and *Ã* within bootstrap samples to quantify how well the description using a single lag thread fits the full matrix of lags (Fig. 2e). To estimate the frequency-dependency of lag threads, we performed a principal component analysis of the primary lag threads across frequency bands (Fig. 2f).

### Relating lag threads to subject age and handedness

To compare subject handedness to lag threads in different frequency bands, we correlated the average EHI handedness score ^22^ with a score for bilateral asymmetry of lag threads across bootstrap samples (Fig. 3c). For each cortical source, we defined the bilaterally antisymmetrized lag thread as the difference of the lag thread and its value at the contralaterally matched cortical source. To gauge the statistical significance of these correlations, we averaged the absolute value of the correlation across the cortex and computed the same statistic for 1000 permutations of the associations between subjects and handedness scores (Fig. 3a). We used Benjamini-Hochberg false discovery correction ^45^ to estimate the statistical significance across frequency bands.

We computed leave-one out pseudovalues of the average lag between sources for each frequency pair and subject, to visualize the relation between lag thread bilateral asymmetry and handedness across subjects. For the pseudovalue computation, we computed the peak lag *l*_1_ of the average cross-correlation across all 89 HCP subjects, and the peak lag *l*_2,i_ of the average across all subjects except subject i. The pseudovalue of the peak lag is then computed as

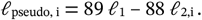

For the correlation between subject age and lag threads, we computed the correlation directly on the pseudovalue level. We defined the time scale of the lag thread for a given subject and frequency as the root mean square (RMS) of the primary lag thread. We then computed the Pearson correlation between the frequency-specific time scale and subject age (Fig. 3d-f).

To assess the statistical significance of the observed correlation, we compared the measure correlation with the distribution of correlations of a null model with permuted association between subject’s MEG data and chronological age. We used false discovery rate correction 45 to control for multiple comparisons across frequencies.

To determine the spatial localization of the correlation effect (Figure 3f), we computed the z-score of lag thread values at each given frequency and cortical location across subjects. The resulting value will thus be larger in areas where the longer temporal scale of the subject-specific lag thread has the largest effect. We then computed the inner product of these values with the z-score of subject ages to obtain the frequency- and location-specific correlation between age and lag thread. We compared this correlation with the corresponding quantity of the null model and performed FDR correction across space to obtain a map of significant clusters of lag thread-age association (Fig. 3f).

### fMRI lag threads

To compare lag threads between MEG and fMRI, we used preprocessed resting state fMRI data from the HCP project for the same subjects whose MEG data we analyzed. We resampled the cortical time series provided by HCP to the 457-point MEG source model by setting the time course at each cortical source to be the weighted average of the HCP preprocessed fMRI time series at the neighboring cortical voxels. Here, the weight of a voxel was proportional to a spatial Gaussian distribution with a standard deviation of *σ* = 5 mm.

We computed the cross-correlogram for each pair of cortical sources for lags ranging between 0 and 20 TRs (corresponding to 14.4 s). We averaged the cross-correlograms using the same bootstrap samples as for the MEG data and computed the lag with the highest cross-correlation using the same algorithm as for the MEG amplitude coupling cross-correlations. We computed the primary lag thread in fMRI by averaging the resulting lag over all outgoing connections (Fig. 4a).

### Comparison between MEG and fMRI lag threads

To compute the similarity between MEG and fMRI threads, we correlated the fMRI lag thread with the MEG lag threads of each frequency and bootstrap sample (Fig. 4b). To test whether subject-specific variation was shared between fMRI and MEG lag threads, we fitted a linear ridge regression, with λ = 10^3^ to prevent overfitting, where the dependent variable was the lag thread in fMRI and the regressors were the MEG amplitude-correlation lag threads. We fitted this model for each combination of bootstrap samples for fMRI and MEG lags. Thus, the bootstrap samples in MEG and fMRI could either be matched or different.

To determine the subject specificity of this linear fit, we correlated the predicted and observed fMRI lag threads to determine how well the fMRI lag thread could be modelled by a linear combination of MEG lag threads. Our hypothesis was that the fit would be better when the bootstrap samples used for the fMRI and MEG lag threads contained the same average over subjects than when they were independently resampled. Accordingly, we computed the difference between the matching and nonmatching linear models. We obtained a positive matched-sample advantage of about 0.8%, indicating that models fit better when MEG and fMRI lag threads were derived from the same subject samples.

To estimate statistical significance, we re-ran the analysis 1000 times with permuted assignments of fMRI and MEG bootstrap samples, which allowed us to estimate whether the better fit between MEG and fMRI lag threads in subject-matched bootstrap samples was statistically significant.

### Cognitive tasks

To compare lag threads across experimental conditions, we first averaged the cross-correlation across sessions using 100 subject-matched bootstrap samples between rest and task, both for the HCP working-memory data and for the locally recorded memory-recall data. We then computed the lag maxima and primary lag threads as described above. We computed the correlation between the primary lag threads in rest and task within each bootstrap sample to estimate the similarity between lag threads in rest and task conditions (Figs. 5a and 6a). We tested for a significant difference by performing a one-sample t-test of the mean from zero using the bootstrap standard error. We used FDR correction to correct for multiple comparisons across frequencies (horizontal bars in Figs. 5a and 6a). For the memory recall task, we additionally computed the hemisphere-wise average of the primary lag threads. This difference corresponds to the average lag between reconstructed sources in the left and right hemisphere.

We compared the mean difference within each frequency band and bootstrap sample between rest and task against the distribution of differences obtained by randomly permuting rest and task data across subjects (Figs. 5a and 6b). We visualized the task effect on lag threads by subtracting rest lag-threads from task lag-threads (Fig. 5b right column; Fig. 6c).

## Footnotes

1 Here, we use a complex-valued continuous wavelet transform using a Morlet wavelet. The method generalizes to any other symmetric, complex-valued wavelet or a short-time Fourier transform (STFT).

